# TGFβ signaling curbs cell fusion and muscle regeneration

**DOI:** 10.1101/557009

**Authors:** Francesco Girardi, Anissa Taleb, Lorenzo Giordani, Bruno Cadot, Asiman Datye, Majid Ebrahimi, Dilani G. Gamage, Douglas P. Millay, Penney M Gilbert, Fabien Le Grand

## Abstract

Fusion of muscle progenitor cells is necessary for skeletal muscle development and repair. Cell fusion is a multistep process involving cell migration, adhesion, membrane remodeling and actin-nucleation pathways to generate multinucleated myotubes. While the cellular and molecular mechanisms promoting muscle cell fusion have been intensely investigated in recent years, molecular brakes restraining cell–cell fusion events to control syncytia formation have remained elusive. Here, we show that transforming growth factor beta (TGFβ) signaling is active in adult muscle cells throughout the fusion process and reduce muscle cell fusion independently of the differentiation step. In contrast, inhibition of TGFβ signaling enhances cell fusion and promotes branching between myotubes. Pharmacological modulation of the pathway *in vivo* perturbs muscle regeneration after injury. Exogenous addition of TGFβ protein results in a loss of muscle function while inhibition of the TGFβ pathway induces the formation of giant myofibres. Transcriptome analyses and functional assays revealed that TGFβ acts on actin dynamics and reduce cell spreading through modulation of actin-based protrusions. Together our results reveal a signaling pathway that limits mammalian myoblast fusion and add a new level of understanding to the molecular regulation of myogenesis.

## Introduction

The adult skeletal muscle cell is a syncytial myofibre that contains hundreds of myonuclei. Formation and regeneration of the myofibre requires fusion of mononuclear progenitors (myoblasts) to form multinucleated myotubes. Located in a niche around the myofibres are quiescent muscle stem cells ^1^, called satellite cells, which can activate and proliferate to give rise to adult myoblasts competent to fuse with each other and with myofibres ^2^. As such, cell fusion plays essential roles in the adult, allowing physiological muscle hypertrophy ^3,4^ and muscle regeneration following injury ^5,6^.

When induced to fuse, adult myoblasts exit the cell cycle, commit to terminal differentiation and migrate toward each other ^7^ They then adhere through membrane integrins ^8^ and cadherins ^9^. The later stages of fusion are controlled by the muscle-specific protein Myomaker ^10^ and peptide Myomerger ^11,12,13^ (also known as Minion, Myomixer). Together, Myomaker and Myomerger reconstitutes cell fusion. Recent studies demonstrated that muscle cell fusion is promoted by actin-based structures ^14^ generating protrusive forces ^15^ and membrane stress then coalescence ^16^. The fusion process must be tightly controlled to safeguard that fusogenic myoblasts do not form aberrant hypertrophic syncytia or fuse with non-muscle cells. However, while it is known that muscle cell fusion can be prevented by tetraspanins at the cell membrane ^17^, no signaling pathway that can limit this process and prevent unscheduled cell fusion has been identified.

Canonical Wnt/β-catenin signaling is a crucial regulator of satellite cells and adult muscle regeneration ^18,19^. Interestingly, β-catenin activation in muscle precursor cells induces the expression of TGFβ ligands and receptors ^20^. TGFβ signaling has been shown, mainly *in vitro*, to negatively regulate myoblast differentiation through functional repression of the myogenic regulatory factors Myod1 ^21^ and Myogenin ^22^. However, TGFβ signaling has broader function in muscle cells, including quiescence ^23^ and activation ^24^ and the impact of TGFβ in syncytia formation has never been investigated. This lack in our knowledge mainly comes from the fact that the primary effects of TGFβ over-expression in skeletal muscles are development of endomysial fibrosis, due to its role as a potent growth factor for connective tissue cells ^25,26^. Here, we asked whether TGFβ signaling might have a role in myoblast fusion.

## Results

TGFβ ligands bind to TGFBR2 that will recruit a type I receptor dimer. The receptor complex will then phosphorylate SMAD2 and SMAD3 that will accumulate in the nucleus where they act as transcription factors ^27^. To evaluate TGFβ isoforms expression by adult muscle progenitor cells, we purified limb muscle satellite cells and grew them *in vitro* as primary myoblasts. We observed that Tgfbl and Tgfb2 expression levels were high in proliferating cells, and diminished following induction of differentiation, while Tgfb3 expression pattern showed an opposite trend (Fig. 1a). Of note, the expression levels of the TGFβ receptors (Tgfbrl, also known as Alk5; and Tgfbr2) did not significantly change during the course of *in vitro* myogenesis (Fig. 1b). We next investigated the state of TGFβ signaling in primary myoblasts, differentiated myocytes and multinucleated myotubes. We observed that the expression level of the TGFβ/SMAD2/3 target gene Smad7 diminished during myogenic progression (Fig. 1c). While immunolocalization of phosphorylated-SMAD2/3 proteins showed that the canonical TGFβ pathway is active at all studied stages (Fig. 1e), quantitative western blotting experiments demonstrated that the intensity of TGFβ signaling decreases during muscle cell differentiation but is not abrogated in multinucleated cells (Fig. 1d). Previous work has established that TGFβ ligands are secreted during muscle tissue repair ^28^. Gene expression analysis of regenerating *Tibialis Anterior* (TA) muscles demonstrated that the 3 TGFβ isoforms are dynamically expressed following injury (Fig. 1e) and peak between 3- and 5-days post-injury (d.p.i.). Likewise, we detected the expression of phosphorylated SMAD3 proteins in nuclei both inside and outside the regenerating myofibres at 4 d.p.i (Fig. 1f). We thus aimed to investigate the role of TGFβ in the fusion process.

**Figure 1.**
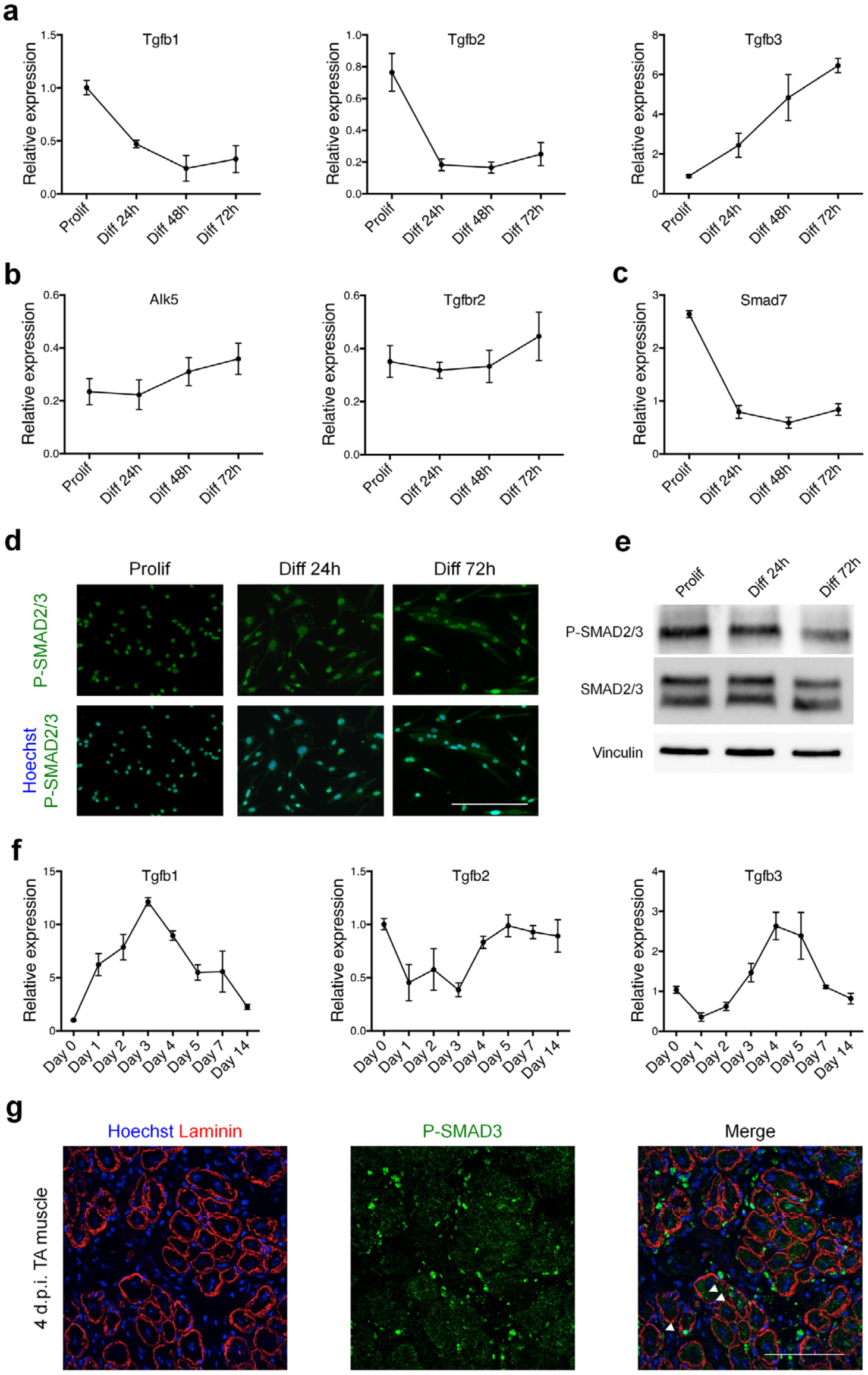
TGFβ signaling pathway remains active during myoblast differentiation. **a,** qRT-PCR analysis of Tgfb1, 2 and 3 transcripts expression during *in vitro* differentiation of primary muscle cells shows different profiles. **b,** qRT-PCR analysis of Alk5 and Tgfbr2 transcripts expression describes a constant expression of the receptors during primary muscle cells differentiation. **c,** qRT-PCR analysis of the TGFβ target gene Smad7 transcript expression reveals a decreased activity of the pathway alongside *in vitro* primary muscle cell differentiation. **d,** Phospho-SMAD2/3 immunofluorescent staining of proliferating, differentiating and differentiated primary myoblasts reveals a constant and basal activation of the pathway. **e,** Phospho-SMAD2/3 and SMAD2/3 western-blot analysis of proliferating, differentiating and differentiated primary myoblasts confirms a decrease in SMAD2/3 phosphorylation during differentiation. **f,** qRT-PCR analysis of Tgfb1, 2 and 3 transcripts expression during muscle tissue regeneration induced by CTX injection shows specific expression profiles. **g,** Immunofluorescent staining for phospho-SMAD3 on 4-days regenerating TA muscles cryosections confirms the presence of active TGFβ signaling in the tissue. Arrowheads indicate phospho-SMAD3^+^ nuclei within myofibres. Scale bars: **e,** 200μm. **g,** 200μm. Data are presented as mean ± s.e.m. from at least three independent experiments.

Since all 3 TGFβ ligands are expressed by cultured satellite cells, we evaluated the impact of recombinant proteins stimulations on adult myogenesis (Fig. S1a). After 72 hours of differentiation, muscle cells aggregated to form multinucleated myotubes, while addition of recombinant TGFβ proteins forced muscle cells to remain mostly mononucleated (Fig. S1b). Quantification of *Myh3* gene expression, which codes for the embryonic myosin heavy chain isoform, further indicated that the cells in TGFβ-treated cultures were in a less mature state than control cultures (Fig. S1c). However, quantification of the percentage of differentiated nuclei expressing pan-MYOSIN HEAVY CHAIN proteins revealed that the vast majority (>90%) of myoblasts did undergo differentiation in all conditions (Fig. S1d), suggesting that TGFβ signaling does not primarily block muscle cell differentiation (Fig. S1e).

To test this hypothesis, we adapted the protocol used by Latroche and colleagues to uncouple differentiation and fusion of primary muscle cells ^29^. In this experimental set up, primary myoblasts were differentiated for 2 days at low density that does not allow contact between cells, split and re-plated at a high density, and cultured for two more days to evaluate muscle cell fusion (Fig. 2a). Following re-plating, almost all muscle cells were terminally differentiated and expressed MYOGENIN (>94%) (Fig. 2b). We thus evaluated the effect of TGFβ proteins stimulations in this experimental setting, *i.e.* on mononucleated differentiated muscle cells (myocytes) and observed that all 3 TGFβ isoforms strongly inhibited cell fusion (Fig. 2c) while muscle cells remained differentiated (Fig. 2d). Activation of the TGFβ pathway reduced the fusion index (Fig. 2e) and completely blocked the formation of large myotubes (Fig. 2f), thus demonstrating that TGFβ signaling limits muscle cell fusion independently of myogenic differentiation. Importantly, addition of TGFβ proteins to adult muscle progenitor cells did not alter their proliferation (Fig. S2a), nor did it induce programmed cell death (Fig. S2b), and it did not alter their motility (Fig. S2c).

**Figure 2.**
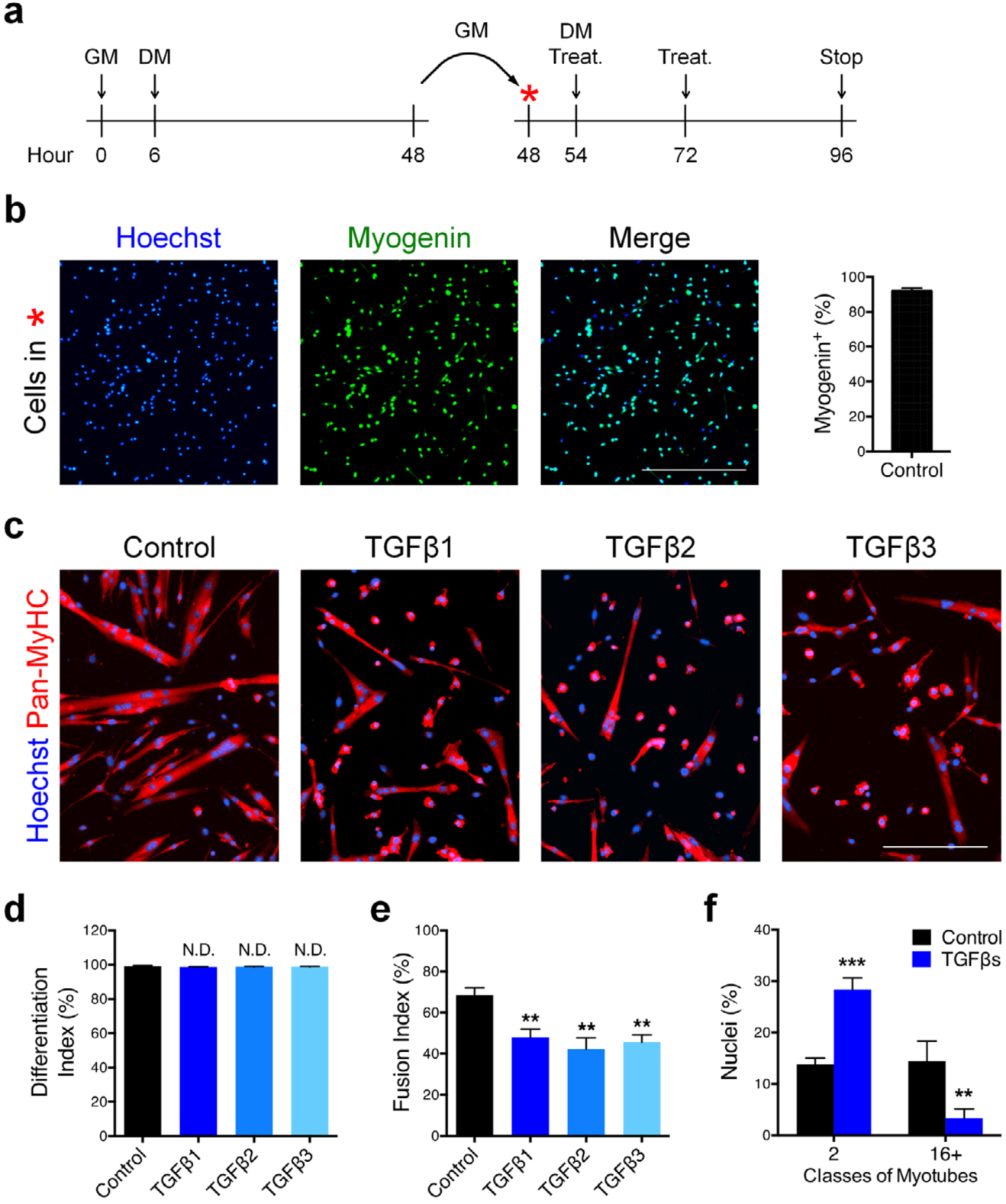
TGFβ signaling limits cell fusion. **a,** Experimental scheme. Primary myoblasts seeded at low density (5,000 cells/cm2) were differentiated for two days, split and re-plated at high density (75,000 cells/cm2) and cultured for two more days. **b,** Immunofluorescent staining for MYOGENIN of primary muscle cells pre-differentiated for 48h and re-plated at high density, confirms that more than 90% of cells express Myogenin. **c,** Immunofluorescent staining for the Myosin Heavy Chain isoforms (Pan-MyHC) of re-plated primary myotubes cultured for 48 hours. **d,** Percentage of Pan-MyHC-expressing cells of re-plated myotubes shows that cells were differentiated in all conditions. **e,** Fusion index of re-plated myotubes reveals that TGFβ stimulation inhibits fusion. **f,** Percentage of nuclei in the smallest and largest myotube classes. TGFβ-treated myotubes are characterized by less nuclei per myotube. Scale bars: **b,** 400μm c, 200μm. Data are presented as mean ± s.e.m. from at least three independent experiments. \*\**P*<0.01, \*\*\**P*<0.001, N.D.=Not significant, compared with Control (Unpaired two-tailed Student’s t-test).

We next asked if inhibition of TGFβ signaling in fusing muscle cells could enhance the formation of multinucleated myotubes. To this aim, we selected ITD-1, a highly selective TGFβ inhibitor which triggers proteasomal degradation of TGFBR2 ^30^. ITD-1 clears TGFBR2 from the cell surface and selectively inhibits intracellular signaling. ITD-1 treatment of primary myocytes resulted in reduced expression of TGFβ target genes (Fig. 3a) and blocked the phosphorylation of nuclear SMAD2/3 proteins induced by TGFβ1 treatment (Fig. 3b). Importantly, treatment of differentiated mononucleated muscle cells re-plated at high density (as in Fig. 2a) with ITD-1 enhanced the fusion process (Fig. 3c). As such, ITD-1-treated cultures showed higher fusion index (Fig. 3e) and were composed of myotubes containing more nuclei (Fig. 3d), of larger diameter (Fig. 3f) and characterized by an aberrant branched shape (Fig. 3g). Taken together our data demonstrate that inhibition of TGFβ receptor function leads to improved fusion and suggest that the levels of TGFβ signaling must be tightly controlled to ensure proper syncytia formation.

**Figure 3.**
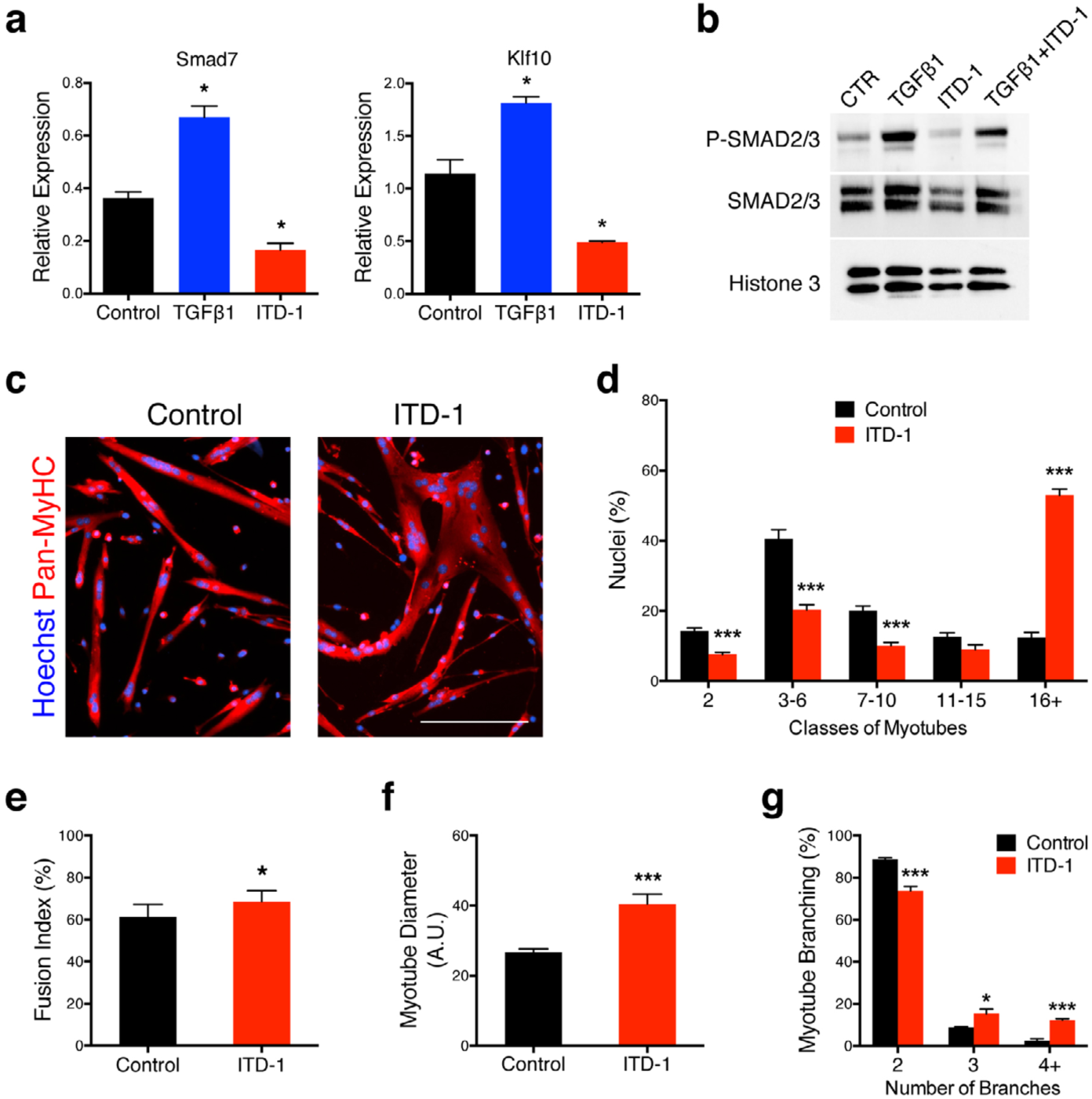
Inhibition of TGFBR2 function in differentiated muscle cell enhance fusion. **a,** qRT-PCR analysis of TGFβ target genes transcript expression in primary myocites treated with TGFB1 protein or ITD-1 compound proves that Smad7 and Klfl0 are over-expressed when the signaling pathway is activated and inhibited when TGFβ cascade is blocked, **b,** Nuclear phospho-SMAD2/3 and SMAD2/3 western blot analysis of primary myoblast treated with TGFB1 protein, ITD-1 compound or both combined. The intracellular mediators SMAD2/3 are phosphorylated upon TGFβ stimulation, while ITD-1 is able to reduce their phosphorylation. **c,** Immunofluorescent staining for Pan-MyHC of re-plated primary myotubes cultured for 48 hours. **d,** Aggregation index of re-plated myotubes shows that ITD-1 treatment leads to the formation of myotubes with higher numbers of nuclei compared to the control. **e,** Fusion index of re-plated myotubes confirms the enhanced fusion when TGFβ cascade is inhibited. Quatntification of the diameter of re-plated myotubes (**f**) and of the distribution of branched-myotubes (**g**) of re-plated cells highlight aberrant morphology of syncitia treated with ITD-1. Scale bars: **c,** 200μm. Data are presented as mean ± s.e.m. from at least three independent experiments. \**P*<0.05, \*\*\**P*<0.Q01, compared with Control (Unpaired two-tailed Student’s t-test).

To test if TGFβ signaling controls fusion cell-autonomously, we expanded primary myoblasts from satellite cells expressing either H2B-GFP or membrane tdTomato (pseudocoloured in blue). Both primary cell types were pre-differentiated at low density, but only the GFP-expressing myoblasts were treated with either TGFβ1 or ITD-1 and their F-actin content stained with SiR-Actin (pseudocoloured in red) (Fig. 4a). Cells were then mixed, replated at high density, and fusion events were imaged live (Fig. 4b). We observed that fusion of GFP-labeled myonuclei into tdTomato myotubes was controlled by the intrinsic state of TGFβ signaling in the fusing cells (Fig. 4c). As such the rate of heterologous fusion was controlled by TGFβ signaling (Fig. 4d). Interestingly, we noticed that multinucleated myotubes could fuse together and that TGFβ signaling regulates the frequency of myotube-to-myotube fusion (Fig. 4e). These results suggest that TGFβ acts cell-autonomously to limit the fusion between muscle cells, and to prevent fusion between syncytia.

**Figure 4.**
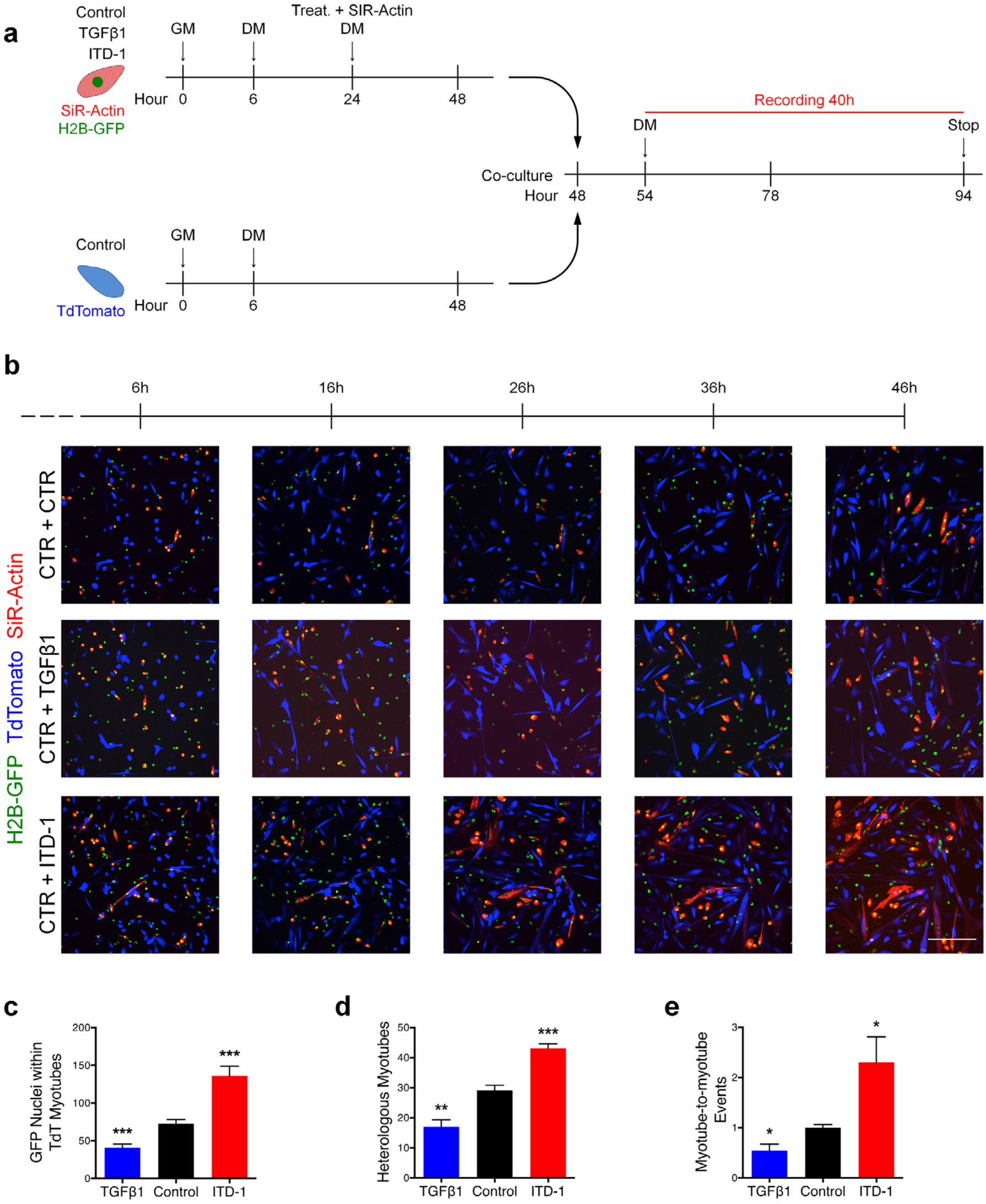
Live-imaging of myoblast fusion. **a,** Experimental scheme. H2B-GFP primary myoblasts were seeded at low density (5000 cells per cm2), treated with TGFβ1 protein or ITD-1 compound, stained with SiR-Actin and differentiated for 2 days. Membrane-TdTomato primary myoblasts were seeded at low density (5000 cells per cm^2^) and differentiated for two days. Both populations were split and co-cultured (50/50) at high density (50,000 cells/cm^2^) and cultured for two more days. In the last 40 hours, cells were recorded live by confocal microscopy. **b,** Live-imaging frames of co-cultured pre-differentiated myoblasts confirm the phenotype previously observed. TGFβ activation inhibits fusion, ITD-1 enhance fusion. **c,** Quantification of H2B-GFP nuclei within TdTomato myotubes. **d,** Quantification of heterologous myotubes (double positive for SiR-Actin and TdTomato). **e,** Quantification of Myotube-to-Myotube events. ITD-1 treatment allows more myotube-to-myotube events compared to the control. Scale bars: **b,** 200μm. Data are presented as mean ± s.e.m. from at least three independent experiments. \**P*<0.05, \*\**P*<0.01, \*\*\**P*<0.001, compared with Control (Unpaired two-tailed Student’s t-test).

To determine the human relevance of the hyper-fusion phenotype obtained with mouse cells, we decided to evaluate the influence of TGFβ pathway inhibition on human muscle cell differentiation in a 3D format (Fig. 5a). By quantifying the tissue remodeling and compaction process (Fig. 5b, 5c), we found that inhibition of TGFBR1 function by the small molecule SB-431542 did not change the thickness of human muscle microtissues (hMMTs) at early differentiation timepoints (Days 0-4), but that as the hMMTs further matured, there was a bifurcation and hMMTs treated with the TGFBR1 inhibitor were significantly thicker than control hMMTs. In this system, hMMT thickening was due to an increase in the width of individual fibres (Fig. 5d, 5e). Notably, SB-431542-treated human muscle fibres contained more nuclei than control fibres, confirming that increased muscle cell fusion as the underlying cellular mechanism (Fig. 5f). By treating hMMTs with acetylcholine to induce tissue contraction and capturing short videos to visualize the magnitude of vertical rubber post deflections, we found that TGFBR1 inhibition renders the hMMTs stronger than their control counterpart Fig. 5g, 5h). Together these data demonstrate that TGFβ signaling regulates human cell fusion.

**Figure 5.**
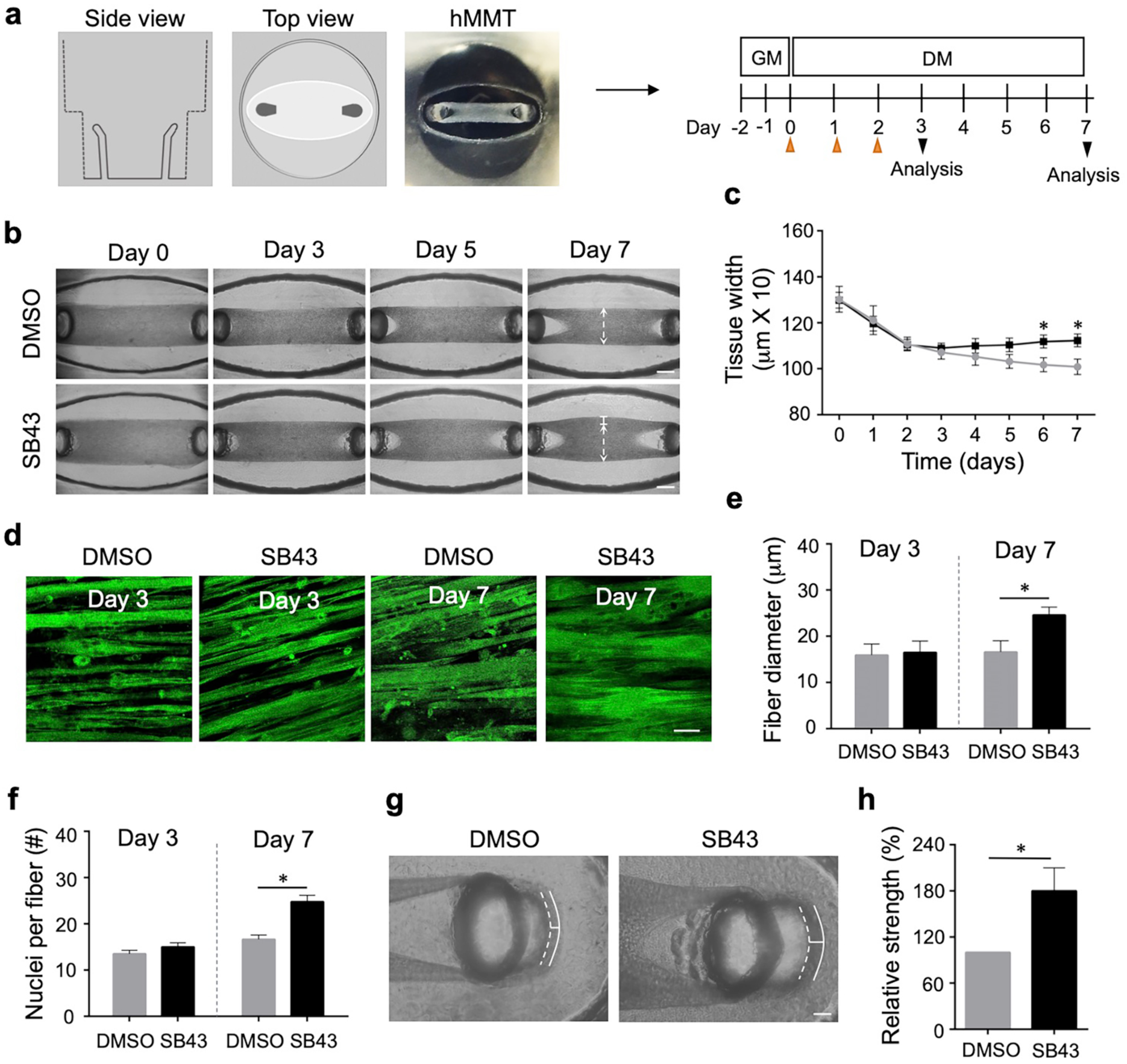
TGFP inhibition induces human myotube fusion in 3D culture resulting in increased microtissue strength. **a,** Schematic representation (left) and timeline (right) of 3D human muscle cell experimental approach utilized in (b-h). Briefly, immortalized human myoblasts are suspended in a fibrin / reconstituted basement membrane protein scaffold and seeded into the bottom of a custom rubber 96-well plate culture device. A side view depicts the vertical posts across which the cells remodel the protein scaffold, align, and fuse to form a 3D human muscle microtissue (hMMT). For the first two days of culture (Day −1, Day −2), tissues are maintained in growth media (GM). On Day 0, GM is removed from wells and replaced with differentiation media (DM). TGFBR1 inhibitor SB-431542 (SB43, 10 uM) was included in the DM on Day 0 – Day 2 (orange arrowheads) of culture. **b,** Representative bright-field images of 3D hMMT culture over the time-course of differentiation treated with SB43 as compared to DMSO-treated control. White arrows demarcate the region of tissues that are assessed in (c). **c,** Line graph quantifying hMMT width over the time-course of differentiation in DMSO (grey line) or SB43 (black line) conditions. **d,** Representative confocal slices of hMMT cultures immunostained for sarcomeric α-actinin (green) on Days 3 and 7 of culture. **e-f,** Bar graph quantifying muscle fiber diameter (**e**) and average number of nuclei per fiber **(f)** at Days 3 and 7 of culture. **g,** Representative brightfield images of hMMTs. Micro-post position before (solid white line) and after (dashed white line) acetylcholine stimulation is represented. **h,** Bar graph quantifying relative strength of SB43-treated hMMTs compared to DMSO-treated hMMTs. Scale bars: **b,** 500μm, **d,** 50μm, **g,** 100μm. n = 3 biological replicates with at least 2 hMMTs replicates per experiment. A minimum of 30 microscopic images per culture condition was analyzed. Data are presented as mean ± s.e.m.. * P< 0.05 compared with Control (Unpaired two-tailed Student’s t-test).

To evaluate the role of TGFβ signaling in muscle cell fusion *in vivo*, we injured TA muscle of adult mice and injected either TGFβ1 protein or ITD-1 compound, at 3 d.p.i., at the time the fusion starts occurring (Fig. 6a). Evaluation of regenerating tissues at 7 d.p.i. revealed that both treatments lead to striking changes in myofibres size and morphology (Fig. 6b). TGFβ1 addition resulted in a robust decrease in nuclear number in newly-formed myofibres compared with controls (Fig. 6e), resulting in a dramatic drop in fibre cross-sectional area (CSA) (Fig. 6c, 6d). In contrast ITD-1 induced a large increase in myonuclear accretion (Fig. 6e) resulting in the formation of larger fibres compared to controls (Fig. 6c, 6d). To further elucidate if modulating TGFβ signaling *in vivo* affects regenerated muscle tissue structure and function, we performed TA muscle injury, followed by 3 successive injections of TGFβ1 or ITD-1, and evaluated the regenerated muscles 2 weeks after injury (Fig. 6f). In this setup, the effects of modifying TGFβ signaling were more pronounced (Fig. 6g). Activation of TGFβ signaling induced the formation of very small myofibres while ITD-1 treatment generated giant myofibres (Fig. 6h, 6i). We next performed *in situ* force measurement of regenerated TA muscles. As suggested by the severe myofibre atrophy observed in TGFβ1-injected muscles, ectopic activation of TGFβ signaling during tissue regeneration lead to a strong reduction of muscle specific force (Fig. 6j). Surprisingly, while being composed of larger myofibres, compared to control regenerated muscles, ITD-1-treated muscles did not show any improvement in force generation, suggesting that increased fibre size resulting from enhanced fusion is not beneficial to muscle function. These observations indicate that TGFβ signaling determine the numbers of fusion events occurring during tissue regeneration *in vivo.*

**Figure 6.**
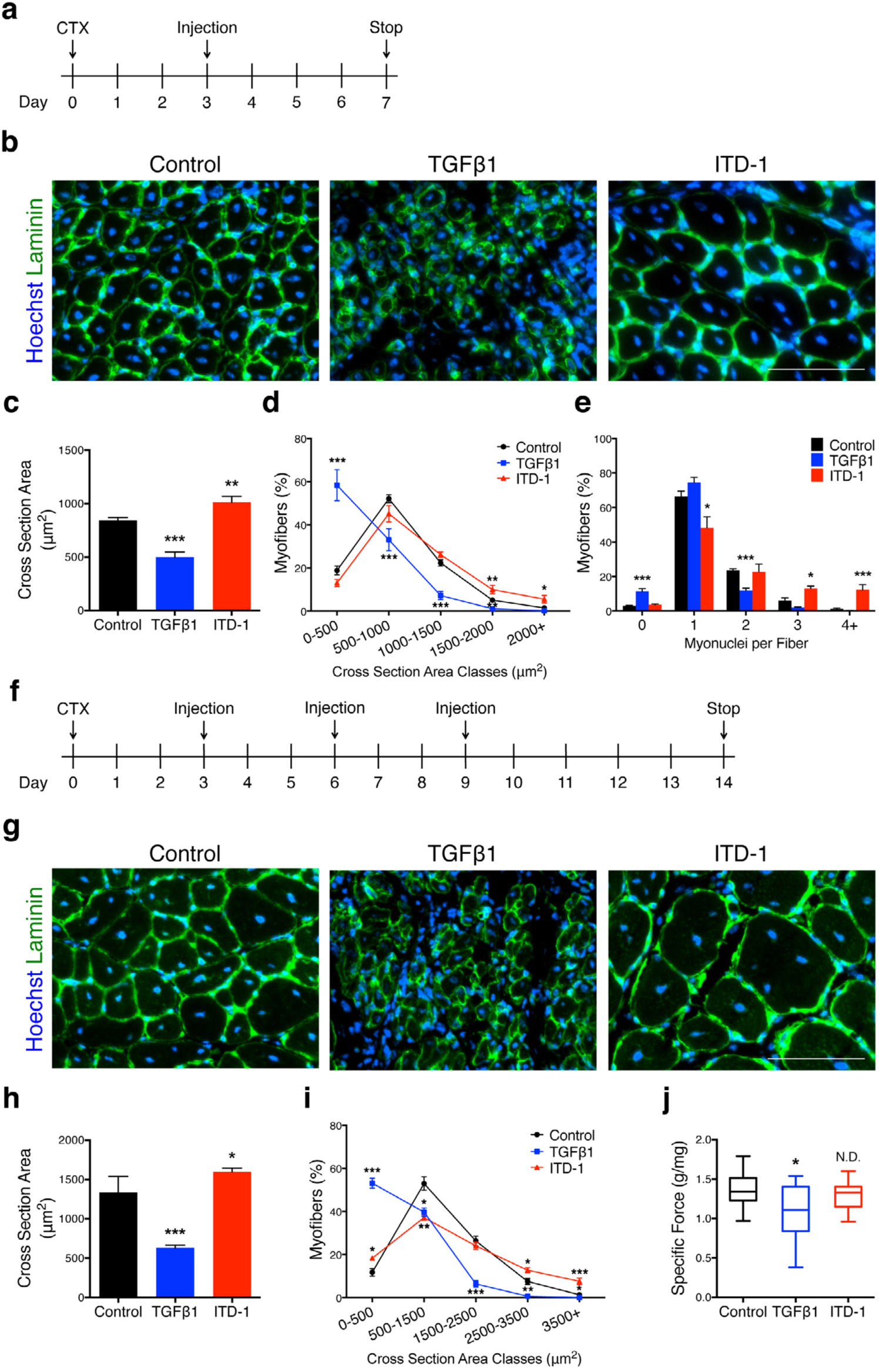
TGFβ signaling regulates muscle cell fusion *in vivo*. **a,** Experimental scheme. Adult mice TA muscles were subjected to CTX injury and regenerating tissues were injected with either TGFβl proteins or ITD-1 compound 3 days after damage. **b,** Immunofluorescent staining for LAMININ of 7-days regenerating TA muscles. **c,** Quantification of myofiber size (cross sectional area, CSA). While the injection of TGFβ strongly reduces the fibers size, ITD-1 administration increases the CSA. **d,** Distribution of myofiber CSA. **e,** Distribution of myonuclei per fiber shows that the inhibition of TGFβ cascade leads to the formation of multi nucleated myofibers, while TGFβ activation reduces the number of myonuclei per fibers. **f,** Experimental scheme. Adult mice TA muscles were subjected to CTX injury and regenerating tissues were injected with either TGFβ proteins or ITD-1 compound 3, 6 and 9 days after damage. 14 days after injury, force measurements were performed and TA muscles collected. **g,** Immunofluorescent staining for LAMININ of 14-days regenerating TA muscles. **h,** Quantification of myofiber size confirms the phenotypes observed at 7d.p.i.. **i,** Distribution of myofiber CSA. **j,** Specific force measurement of regenerating muscles. While TGFβ1-treated muscles are weaker compare to the control, ITD-1 injected muscles are funstional. Scale bars: **b, g**, 100μm. Data are presented as mean ± s.e.m. from at least three independent experiments. \**P*<0.05, \*\**P*<0.01, \*\*\**P*<0.001, N.D.=Not significant, compared with Control (Unpaired two-tailed Student’s t-test). Control represents mock-treated contralateral tibialis anterior muscle.

To identify the genetic networks regulated by TGFβ signaling in muscle cells, we performed transcriptome analysis of primary myocytes differentiated for 24 hours and stimulated with either TGFβ1 or ITD-1 for another 24 hours (Fig. 7a). We first observed that the relative expression levels of myogenic transcription factors (Pax7; Myod1; Myogenin) were not significantly changed, confirming that the modulation of TGFβ signaling does not act on myogenesis in these experimental conditions. Recently identified fusion master regulatory factors (Myomaker and Myomixer) were also unaffected by TGFβ signaling (Fig. 7b). We then used Ingenuity Pathway Analysis (IPA) to reveal the pathways affected by TGFβ signaling. Interestingly, we found that “Actin Cytoskeleton” was among the top-regulated pathways (Fig. 7c). This is significant, since actin remodeling and the formation of finger-like actin protrusions are essential for myoblast fusion ^31^. IPA further revealed changes in the transcription of numerous genes implicated in actin dynamics following TGFβ treatment (Fig. S3a). Visualization of the F-ACTIN network and measure of the local orientation of actin filaments in differentiated muscle cells showed that the level of TGFβ signaling negatively correlates with cytoskeleton reorganization (Fig. 7d) and elongated cell shape with an effect on cell spreading (Fig. 7e) and the coherency of actin filament alignment (Fig. 7f).

**Figure 7.**
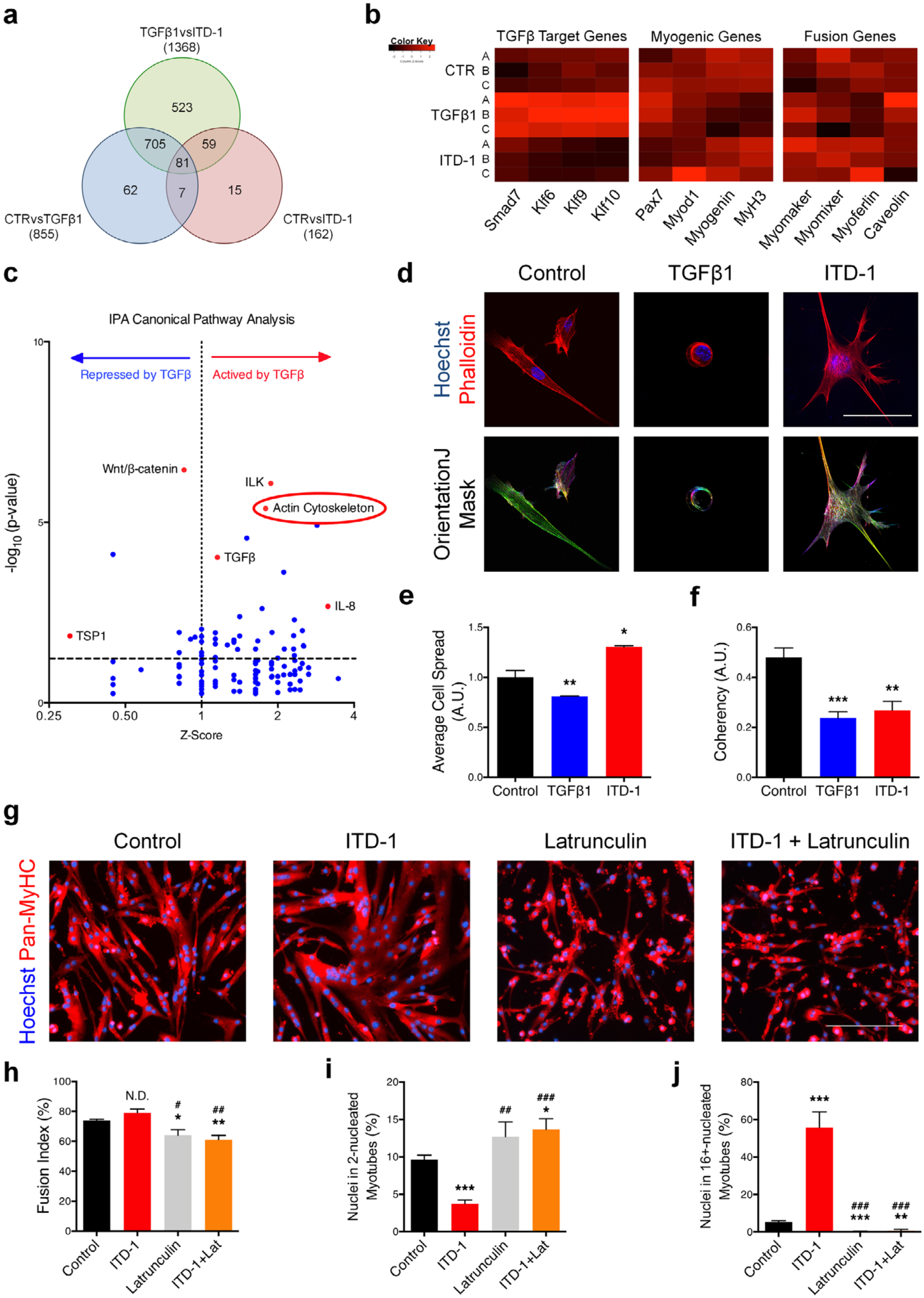
Fusogenic actin remodeling is controlled by TGFβ signaling. Transcriptomic analysis was performed on differentiated myocytes treated with either TGFBI or ITD1. **a,** Venn Diagram showing differentially expressed gene overlap across the three conditions. **b,** Heatmaps of TGFβ Target genes, myogenic genes and fusion genes. **c,** Volcano Plot showing the Ingenuity Pathway Analysis (IPA). Among the top modulated pathways, Actin Signaling Pathway is highlighted. **d,** Phalloidin staining of 1-day differentiated myocytes. These pictures were analyzed with OrientationJ (ImageJ Plug-in) to obtain a color-coded orientation mask. **e,** Average cell spread quantification. TGFpi treatment reduces cell size; ITD-1 promotes cell spreading. **f,** Quantification of orientation coherency of the Actin fibers. Both treatments reduce coherency compare to the control. **g,** Immunofluorescent staining for all the Mysosin Heavy Chain isoforms (Pan-MyHC) of re-plated primary myotubes cultured for 48 hours with ITD-1, Latrunculin, or both. **h,** Fusion index of re-plated myotubes shows that Latrunculin significantly reduces the parameter when administrated. **i,** Percentage of nuclei in the smallest myotube classes. ITD-1-treated myotubes are characterized by less nuclei per myotube, while Latrunculin increases the percentage of nuclei in small myotubes when administrated alone or together with ITD-1. **j,** Percentage of nuclei in the biggest myotube classes. ITD-1 strongly increases the number of nuclei in big myotubes, but Latrunculin blunts this effect, reducing the percentage. Scale bars: **d,** 40μm. g, 200μm. Data are presented as mean ± s.e.m. from at least three independent experiments. Coherency was calculated from at least 150 cells per condition. \**P*<0.05, \*\**P*<0.01, \*\*\**P*<0.001, compared with Control. #*P*<0.05, ##*P*<0.01, ###*P*<0.001, compared with ITD-1 (Unpaired two-tailed Student’s t-test).

To better understand actin remodeling during muscle cell fusion, we performed live-imaging experiments to visualize accumulation of F-ACTIN foci in invasive podosome-like structures at the sites of fusion between cells. By mixing cells stained with SiR-Actin and unstained cells, we observed dynamic actin remodeling in untreated cells (Fig. S3b, Top) and cells stimulated with ITD-1 (Fig. S3b, Bottom), while TGFβ1 stimulation prevented the formation of actin-rich invasive structures and promoted the maintenance of a rounded cell shape (Fig. S3b, middle).

Lastly, we asked whether TGFβ-driven effect on fusion is conserved in a non-muscle context. To do so, we took advantage of a fibroblast dox-inducible cell fusion reconstitution system ^32^. Myomaker- and Myomerger-transduced 10T1/2 fibroblasts were seeded and stimulated with dox to induce fusion (Fig. S4a). TGFβ1 recombinant protein was administrated at multiple time points (from day 0 to day 3), but none of the different setting led to a reduction of fusion, compared to the untreated fibroblasts (Fig. S4b, S4c). These results show that TGFβ1 protein is unable to inhibit the fusion process in a non-muscle cell type and that its signaling cascade acts independently from Myomaker and Myomixer, thus confirming the evidences obtained from the transcriptome analysis (Fig. 7b).

## Discussion

The data presented here identify an unexpected negative role for TGFβ in the fusion of adult myoblasts to form myotubes. TGFβ signaling has previously been shown to play a major role in the skeletal muscle morphogenesis. Throughout development, TGFβ ligands are expressed mostly by connective tissue cells and in close proximity to growing muscle tissue ^33^. While it is known that connective tissue cells provide a pre-pattern for limb muscle patterning ^34^ and control the amount of myofibres within the developing muscle masses ^35^, we propose that TGFβ is the main signal limiting muscle cell fusion.

During adult tissue repair TGFβl and TGFβ3 are mainly expressed by inflammatory macrophages invading the regenerating muscle tissue ^36,37^ while TGFβ2 is secreted by activated satellite cells and differentiating myotubes ^36^. This sequential expression of ligands by different cell types may be instrumental in preventing premature fusion between transient amplifying myoblasts and then in avoiding fusion events between syncytia and prevent the formation of aberrant branched myotubes. Myofibres with branches are found in muscular dystrophy ^38^. They present morphological malformations, as well as alterations in calcium signaling ^39^, and arise from asynchronous myofibre remodeling. Knowledge of the signaling pathways regulating muscle cell fusion may help design therapeutic strategies to decrease myofibre branching in dystrophic patients.

Our *in vivo* experiments indicate that TGFβ signaling must be tightly regulated in muscle progenitor cells during tissue repair. Treatment of regenerating muscle tissues with TGFβl protein strongly blocks muscle progenitor cell fusion, and the impair the function of the regenerated tissue. Interestingly, ITD-l administration lead to the formation of giant myofibres containing more nuclei. However, while functional, the regenerated muscle did not generate higher force compared to mock-treated regenerated tissues. While appealing, it remains to be demonstrated if enhancing cell fusion might improve regenerated muscle function. As such, “bigger” does not always means “stronger”, and this is exemplified by our previous analysis showing that lack of Rspol results in the formation of larger myofibres containing supernumerary nuclei following regeneration ^6^.

Work in *Drosophila melanogaster* previously demonstrated that actin polymerization drives muscle cell fusion ^31^. Our demonstration that TGFβ stimulation breaks down actin architecture links extracellular cues to cell mechanics. We show that the state of TGFβ signaling in pre-fusing cells controls their shape, and the formation of actin-based protrusions which are necessary for fusion of mammalian cells ^40^. Our result also demonstrate that blockade of actin polymerization blunts the over-fusion phenotype induced by inhibition of TGFβ signaling. Interestingly, TGFβ stimulation of Myomaker- and Myomerger-expressing fibroblast enhanced fusion of these cells. This can be explained by the fact that TGFβ induces cytoskeletal reorganization and actin remodeling in fibroblasts ^41,42^ in an opposite way as we observed in muscle cells.

In the present state of our knowledge, most of the fusion-promoting factors have been discovered through *in vivo* studies in the fly embryo. As such, the concept of the fusogenic synapse; the site where an attacking fusing cell propels an actin-rich membrane protrusion towards a receiving cells, is poorly characterized in mammalian cells ^43^. Here, we identify numerous actin-related transcripts, of which the expression is regulated by TGFβ signaling, that may be integral parts of the molecular fusion machinery. Further work should thus be dedicated to the study of the TGFβ-regulated genes, and their specific roles in actin dynamics to better our understanding of how muscle cell membranes are brought together for fusion.

In conclusion, the elucidation of TGFβ signaling as a brake for myoblast fusion opens new avenues to study this fundamental cellular process at a molecular level and to understand how fusion is perturbed in neuromuscular diseases.

## Acknowledgement

We thank the CyPS Facility for technical support. We thank C. Coirault and A. Brack, F. Relaix and R. Mounier for commenting the draft manuscript. Work in the F.L.G. lab was supported by grants from the Agence Nationale pour la Recherche (ANR-14-CE11-0026 and ANR-12-JSV2-0003), and from the Association Française contre les Myopathies/AFM Telethon. F.G. was supported by a PhD fellowship from the Fondation pour la Recherche Médicale (ECO20160736081). P.M.G acknowledge the following sources for funding this study: Natural Science & Engineering Research Council (NSERC; RGPIN 435724), NSERC Canada Research Chair’s Program, and the Canada First Research Excellence Fund ‘Medicine by Design’.

## Author Contributions

F.L.G conceived and designed the study. F.G. performed most of the experiments. A.T. performed western blotting experiments. L.G. performed RT-qPCR experiments. F.L.G performed cell culture experiments. B.C. conceived the live imaging experiments. A.D., M.E., and P.M.G. designed, performed, and interpreted human muscle cell culture experiments. D.G.G., and D.P.M. designed, performed, and interpreted the fibroblast experiment. F.G. and F.L.G. wrote the manuscript. All authors reviewed the manuscript.

## Supplementary Information

## Methods

### Mice

Wild-type mice used in this project were 2 to 5 months-old C57Bl6/N mice purchased from Janvier Laboratories. Experiments were performed at the Centre d’Expérimentation Fonctionnelle (UMS28) Animal Facility following the European regulations for animal care and handling. Experimental animal protocols were performed in accordance with the guidelines of the French Veterinary Department and approved by the Sorbonne Université Ethical Committee for Animal Experimentation. Cardiotoxin (CTX) injection in Tibialis Anterior (TA) muscle and hindlimb muscle dissection were performed following the protocol described in ^44^

### Skeletal Muscle Injury

Mice were anaesthetized by intraperitoneal injection of Ketamin at 0,1mg per gram body weight and Xylazin at 0,01mg per gram body weight diluted in saline solution. 30 ul of CTX (12 mM in saline, Latoxan) was injected into hindlimb TA muscles to induce injury, and mice were euthanized 0, 1, 2, 3, 4, 5, 7 or 14 days afterward. Recombinant mouse TGFβ1 (R&D Systems) was diluted in saline and 250 ng (25 ul) was injected into the TA every injection. ITD-1 compound was diluted in saline and 200 ng (25 ul) was injected into the TA every injection. Muscles were freshly frozen in OCT Embedding Matrix compound (CellPath) and cut transversally at 10 um with a Leica cryostat.

### In situ Physiological assay

Tibialis anterior muscles were evaluated by the measurement of in situ isometric muscle contraction in response to nerve stimulation. Mice were anaesthetized intraperitoneal injection of Ketamin at 0,1mg per gram body weight and Xylazin at 0,01mg per gram body weight diluted in saline solution. Feet were fixed with clamps to a platform and knees were immobilized using stainless steel pins. The distal tendons of muscles were attached to an isometric transducer (Harvard Bioscience) using a silk ligature. The sciatic nerves were proximally crushed and distally stimulated by bipolar silver electrode using supramaximal square wave pulses of 0.1ms duration. All data provided by the isometric transducer were recorded and analysed on a microcomputer, using PowerLab system (4SP, AD Instruments). All isometric measurements were made at an initial length L0 (length at which maximal tension was obtained during the tetanus). Responses to tetanic stimulation (pulse frequency from 75 to 143Hz) were successively recorded. Maximal tetanic force was determined. Muscle masses were measured to calculate specific force.

### Murine Cell Cultures

Skeletal muscle-derived primary myoblasts were isolated from wild-type mice using the Satellite Cell Isolation Kit MACS protocol (Miltenyi Biotec). Briefly, hindlimb muscles were dissected out, placed in a sterile Petri dish and minced to a pulp with curved scissor. The pulp was then incubated in a CollagenaseB/DispaseII/CaCl^2^ solution at 37°C for 40 minutes with two trituration steps. The enzymes are then blocked by addition of Fetal Bovine Serum (FBS) and the muscle extract was treated with Red Blood Cell Lysis Solution to remove erythrocytes. After this step, magnetic labeling is performed by adding Buffer (5% BSA, 2mM EDTA, PBS) and Satellite Cell Isolation Kit (a mixture of antibodies specific for non-satellite cells conjugated with magnetic beads). Cell suspension is then poured into the column in the magnetic field. Unlabeled cells (satellite cells) flow through the column while magnetically labeled cells are retained within the column.

Satellite cells were resuspended in growth medium (Ham’s F10 with 20% FBS, 1% penicillin/streptomycin and 2.5ng/ml of bFGF) and plated into a collagen-coated 60mm Petri dish. Cells were maintained in growth medium until cells reached 80% confluence. To induce myogenic differentiation and fusion, myoblasts were plated at different concentrations depending on the experimental design (5000, 20000 or 75000 cells/cm^2^) onto matrigel coated plates in growth medium. Once adherent, cells were incubated in differentiation medium (DMEM with 2% Horse serum and 1% penicillin/streptomycin) for up to 3 days. For recombinant protein treatments, we used TGFβ1 (eBioscience), TGFβ2 (Biotechne) and TGFβ3 (Biotechne) administrated at a final concentration of 20ng/ml. ITD-1 (Tocris) compound was administrated at 5mM.

### PDMS mold fabrication for 3D human muscle microtissues (hMMT)

To generate 3D human muscle microtissues (hMMTs) we employed a second-generation micro-molded device in a 96-well format made of polydimethylsiloxane (PDMS; monomer/cross-linker ratio = 15: 1) in a single simple molding process. At the bottom of each well of the 96-well microfabricated device an oval shaped pool was designed with a vertical flexible PDMS post on each side of it. PDMS culture plates were sterilized via an autoclave. Just prior to use, wells were further sterilized by an overnight incubation with a 5% pluronic acid solution (100μL/well) at 4°C, which also served to create a non-adhesive surface to support tissue self-organization.

### Human skeletal muscle microtissue (hMMT) culture

The 3D hMMTs were generated using human immortalized myoblast lines obtained from V. Mouly (AB1167 from fascia lata muscle of a healthy 20-year old male, AB1190 from paravertebral muscle of a healthy 16-year old male, and KM155 from thigh muscle of a healthy 25-year old male) ^45^. Immortalized myoblasts were cultured in Skeletal Muscle Cell Basal Medium with Skeletal Muscle Cell Growth Medium Supplement Mix (PromoCell) supplemented with 20% FBS and 1% P/S. Myoblasts were harvested by trypsinization and resuspended (1.5 x 10^5^ cells/tissue or 1.0 x 10^7^ cells/mL) in a hydrogel mixture consisting of fibrinogen (4 mg/mL, 40% v/v; Sigma) and Geltrex (20% v/v; Thermo Fisher Scientific) in DMEM (40% v/v) in the absence of thrombin. Then, 0.2 unit of thrombin (Sigma) per each mg of fibrinogen was added just before seeding the cell-hydrogel mixture into the wells and left for 5 min in an incubator at 37°C to allow optimal fibrin gel formation. Subsequently, 200 μL of growth medium consisted of Skeletal Muscle Cell Growth Medium lacking supplement mix (Skeletal Muscle Cell Basal Medium; PromoCell) supplemented with 20% FBS, 1% P/S and 1.5 mg/mL 6-aminocaproic acid (ACA; Sigma) was added to each well. The hMMTs were cultured in the growth medium for two days, and then the medium was replaced to differentiation medium (DMEM supplemented with 2% horse serum, 1% P/S, and 10 μg/mL human recombinant insulin) containing 2 mg/mL ACA to induce differentiation. SB43 (Sigma) was added at the final concentration of 10 μM into the differentiation medium of hMMTs for Day 0 to Day 2 of differentiation, while an equivalent volume of DMSO was added to control samples. On Day 3 of differentiation, a final full media exchange was performed to remove SB43 or DMSO. Half of the culture medium was replaced every other day for the remaining differentiation period.

### 3D tissue compaction and tissue remodelling analysis

The effect of SB-43 treatment on hMMT compaction was evaluated by measuring tissues diameter as an indication for tissue self-organization and remodelling over differentiation period. Phase contrast 4x magnification images were captured over time using an inverted microscope (Olympus) to analyze 3D tissue compaction. In each image four width measurements were done across the length of the tissue using ImageJ software and the average diameter was calculated. The data was shown as absolute diameter change over time and the result was compared between SB43 treated and DMSO treated control tissues.

### hMMT Myofibre width and nuclear index analyses

Myofibre width was measured using 40x magnification stack images of SB43 and DMSO treated hMMTs at Day 3 and 7 day of the differentiation period. Analysis of SAA immunostained images of 3D muscle tissues was facilitated by use of NIH ImageJ software. We analyzed a total of three to five images per tissue, to determine the diameter of each muscle fibre. Myofibres were only qualified for fibre diameter analysis if they were visible across the length of the stacked image. In this work, myotubes were defined as multinucleated cells comprising at least three fused nuclei. Three width measurements were done across the length of each qualified fibre to ensure that the thickest and the thinnest parts were included in the measurements, and subsequently the average fibre diameter per condition was calculated. To determine the average number of nuclei per fibres, we quantified the total number of Hoechsts nuclei contained within each SAA+ muscle fibre in each hMMT culture condition.

### hMMT relative force quantification

To evaluate the effect of TGFβ inhibition on the function of hMMTs, we evaluated hMMTs contraction in response to acetylcholine (ACh; Sigma) stimulation and tissue contraction. Briefly, ACh solution (in DMEM) was directly added (1mM final concentration) into the wells containing hMMTs after 7 days differentiation. We then captured short phase-contrast videos at 10x magnification to visualize the movement of the movement of the flexible PDMS posts. Post displacement was quantified using ImageJ software. Relative hMMTs strength data was evaluated by normalizing the post displacement of SB43 treated tissues to DMSO treated control tissues.

### Myomaker and Myomerger-expressing 10T1/2 fibroblasts

Myomaker and Myomerger-expressing fibroblasts were used and described previously (Leikina et al., 2018). Myomaker and Myomerger transduced 10T1/2 fibroblasts were seeded in 8 - chamber Ibidi slides with a 3 x 10^3^ cell density per well (Day 0). 8 hours of post-seeding (Day 0), Myomerger expression was induced by treating the cells with dox containing culture medium (1 μg/mL) and replaced every 24 hours. Each experimental chamber was treated with human TGFβ1 recombinant protein (20 ng/mL; ebioscience) as specified. 4 days after seeding, cells were fixed, and fusion was evaluated by analyzing the number of nuclei in GFP^+^ cells. The experiment was performed three times in duplicate and at least 3 images per well was quantified.

### RNA Extraction and Quantitative Real Time PCR (RT-qPCR)

Total RNA was isolated from cultured cells and TA muscles using TRIzol Reagent (Thermo Fisher) or Direct-zol RNA Kit (Zymo Research) according to the manufacturer’s protocol. TA muscle tissue was destroyed using the MagNa Lyser System (Roche). RNA concentration was evaluated with Nanodrop. After subsequent DNAse treatment, cDNA was generated using High Capacity Reverse Transcription Kit (Life Technologies). cDNA was then used for quantitative PCR (qPCR) done with LightCycler 480 SYBR Green Master Mix (Roche) and run in LC480 for 40 cycles. Primers are reported in **Table 1**. All samples were duplicated and transcripts levels were normalized for a housekeeping gene relative abundance (TBP, transcription regulator).

**Table 1:**
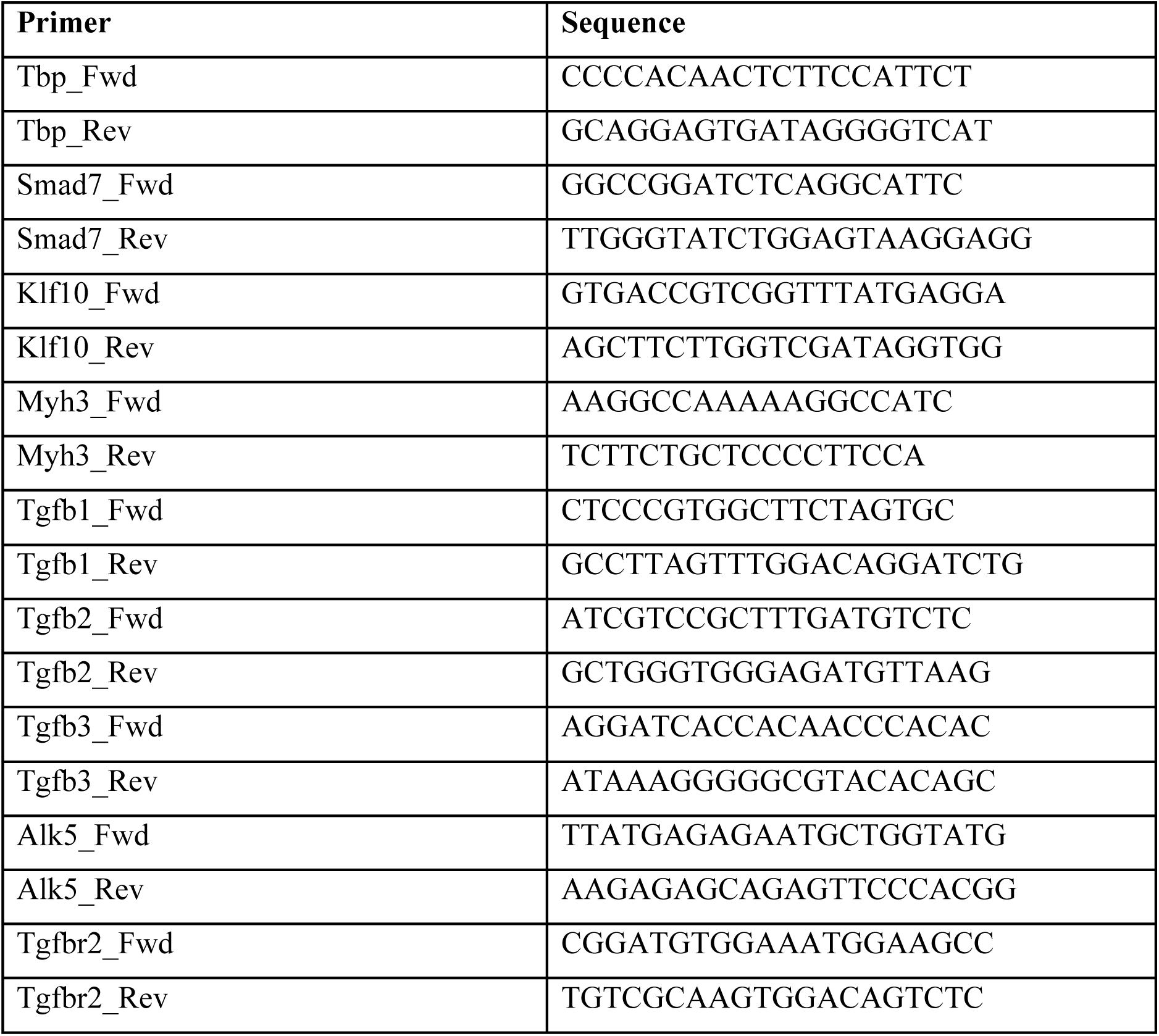
RT-qPCR primers used in this study

### Microarray and bioinformatics

The RNA from primary myoblasts were isolated using Direct-zol RNA Kit (Zymo Research) according to the manufacturer’s protocol. After validation of the RNA quality with Bioanalyzer 2100 (using Agilent RNA6000 nano chip kit), 100ng of total RNA is reverse transcribed following the GeneChip® WT Plus Reagent Kit (Affymetrix). Briefly, the resulting double strand cDNA is used for *in vitro* transcription with T7 RNA polymerase (all these steps are included in the WT cDNA synthesis and amplification kit of Affymetrix). After purification according to Affymetrix protocol, 5.5ug of Sens Target DNA are fragmented and biotin labeled. After control of fragmentation using Bioanalyzer 2100, cDNA is then hybridized to GeneChip® MouseGene2.0ST (Affymetrix) at 45°C for 17 hours.

After overnight hybridization, chips are washed on the fluidic station FS450 following specific protocols (Affymetrix) and scanned using the GCS3000 7G. The scanned images are then analyzed with Expression Console software (Affymetrix) to obtain raw data (cel files) and metrics for Quality Controls. Data were normalized using RMA algorithm in Bioconductor with the custom CDF vs 22. Statistical analysis were carried out with the use of Partek® GS. First, variations in gene expression were analyzed using unsupervised hierarchical clustering and PCA to assess data from technical bias and outlier samples. To find differentially expressed genes, we applied a one-way ANOVA for each gene. Then, we used unadjusted p-value and fold changes to filter and select differentially expressed genes. In TGFβ1 or ITD-1 -treated myocytes, the genes were selected with p<0,05 significance and at least 50% difference. Gene networks and canonical pathways representing key genes were identified using the curated Ingenuity Pathways Analysis ® (IPA) database.

### Muscle Histology and Immunofluorescence

Cell cultures were fixed with 4% paraformaldehyde in PBS for 15 minutes, washed 3 times with PBS and permeabilized with 0.25% Triton-X in PBS for 10 minutes. Blocking step was performed using 4% Bovine Serum Albumin (BSA) for 45 minutes at room temperature. Cultures were then incubated with primary antibodies diluted in blocking solution (4% BSA) overnight at 4°C. After a quick wash with 0.1% Np40 in PBS, samples were washed with PBS twice. Cells were incubated with secondary antibodies for 45 minutes at room temperature. Secondary antibodies used were anti-Mouse IgG or anti-Rabbit IgG coupled with Alexa Fluor 488 or 546 dyes, from Life Technologies. F-Actin staining was performed using FITC-conjugated Phalloidin (Sigma) or SiR-Actin (Spiro Chrome). Following antibody staining, cultures were washed 3 times with PBS and nuclei were stained with Hoechst, before being analyzed with the EVOS FL Cell Imaging System microscope (Life Technologies) or with a Nikon Ti2 microscope equipped with a motorized stage and a Yokogawa CSU-W1 spinning disk head coupled with a Prime 95 sCMOS camera (Photometrics). hMMT samples were fixed with 4% formaldehyde (Alfa Aesar) for 15 min at room temperature (RT). Following three washes with PBS, samples were permeabilized and blocked with a blocking solution containing 10% goat serum (Life Technologies), 0.3% Triton (BioShop) in PBS for 30 min at RT. Samples were then incubated with a mouse anti sarcomeric α-actinin antibody (Sigma) diluted 1:800 in the blocking solution overnight at 4°C. After three washes with PBS, samples were then incubated with goat anti-mouse IgG conjugated with Alexa-Fluor 488 secondary antibody (1:500; Invitrogen) and Hoechst 33342 (1:1000; Invitrogen) diluted in the blocking solution for 60 min at RT. Multiple confocal stacks through each tissue were captured at multiple randomized locations using an Olympus IX83 inverted confocal microscope equipped with FV-10 software.

### Live Imaging

Fluorescent-labeled myoblast cultures were pre-differentiated 48 hours at a low density (5000 cells/cm^2^) onto matrigel coated plates and and re-plated in Nunc Lab-Tek Chamber Slide system at a high density (75000 cells/cm^2^) and cultured for about two more days. Cells were recorded for the last 40 hours of differentiation using a Nikon Ti2 microscope equipped with a motorized stage and a Yokogawa CSU-W1 spinning disk head coupled with a Prime 95 sCMOS camera (Photometrics). Specifically, for each condition and replica 4 fields at 20x magnification were recorded every 10 minutes.

### Western Blot

Cells were lysed with RIPA buffer (Sigma-Aldrich) supplemented with Protease and Phosphatase Inhibitor (Thermo Fisher). If needed, cytoplasmic and nuclear proteins were separated using the NE-PER Nuclear and Cytoplasmic Extraction Reagents (ThermoFisher). Protein quantification was performed using BCA Protein Assay Kit (Thermo Fisher) and after were denatured and reduced incubating the samples with 2X Laemmli Sample Buffer (Santa Crus BioTechnology) for 30 minutes at room temperature. Equal amounts of proteins were loaded in SDS-PAGE gel NuPAGE™ 4-12% Bis-Tris Protein Gels (Thermo fisher) along with molecular weight marker. Load 10μg of total protein from cell lysate and transferred on nitrocellulose membranes (Bio-Rad Laboratories, Inc.). After blocking in 5% milk or BSA and 0.1% Tween-20/TBS, membranes were incubated with primary antibodies (Table 2) overnight and then with HRP-conjugated secondary antibodies for 1 hour. Specific signals were detected with a chemiluminescence system (GE Healthcare).

**Table 2:**
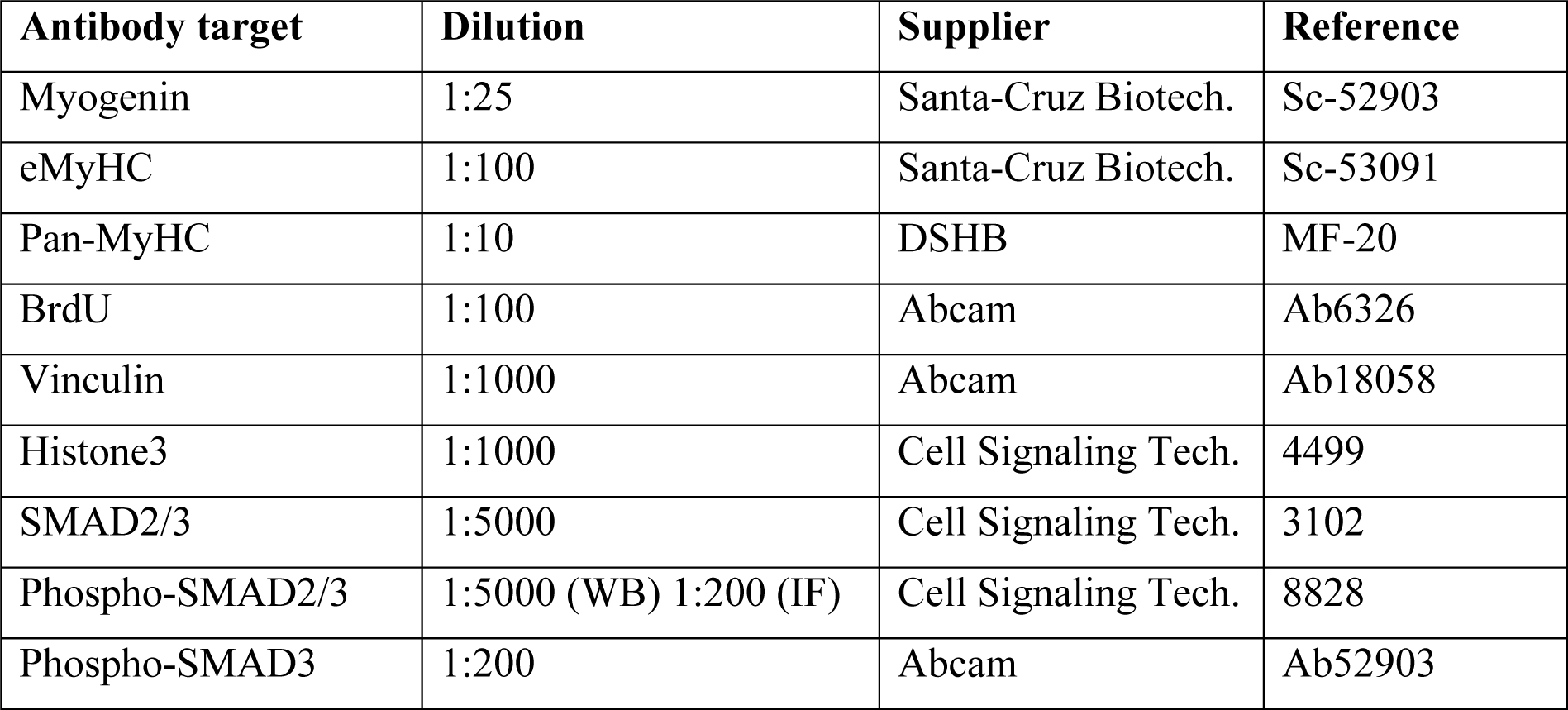
Antibodies used in this study

### BrdU Assay

Cell cultures were grown at 20000 cells per cm^2^ density in growth medium on collagen-coated plates. Cells were treated with TGFβ isoforms for 1 day and then incubated with BrdU for 40 minutes before fixing them with 4% PFA for 5 minutes at RT. After a brief wash in PBS, cells were denaturated with 2M HCl for 30 minutes at 37°C. To neutralize the acid, 6 consecutive washes in PBS of 5 minutes each are performed. Cells were blocked with 2% Goat Serum, 0.2% Tween20 PBS for 30 minutes at 37°C. Cells were incubated with primary antibody (**Table 2**) for 2 hours at RT and then washed 3 times in PBS. Secondary antibody (anti-Rat IgG conjugated with Alexa Flour 546 from Life Technologies) was added and incubated for 45 minutes at RT. Before microscope observation, three washes with PBS and Hoechst staining were performed.

### TUNEL Assay

Cell cultures were grown at 20000 cells per cm^2^ density on collagen-coated plates and treated with TGFβ proteins for 1 day both in proliferating and in differentiating conditions. Cells were fixed in 4% PFA at RT for 20 minutes, washed three times with PBS and permeabilized with 0.1% TritonX-100, 0.1% Sodium Citrate PBS for 2 minutes on ice. After three washes in PBS, cultures were treated according to the protocol of In Situ Cell Death Detection Kit (Roche). Before observation, three washes with PBS and Hoechst staining were performed.

### Scratch-Wound Assay

Primary myoblasts were plated at 30000 cells per cm^2^ density on collagen-coated plates. When cells reached 80% confluence, we scratched the monolayer of cells in a straight line, washed with PBS few times and incubated the cells in growth medium with TGFβ1 or ITD-1. Cells were stained by NucBlue Live ReadyProbes Reagent (Thermo Fisher) to ensure that all cells were removed within the scratch. After 24 hours, myoblasts were fixed with 4% PFA and the number of cells in the scar was counted manually.

### Statistical Analysis

A minimum of 3 biological replicates was performed for the presented experiments. Error bars are standard errors. Statistical significance was assessed by the Student’s t-test, using Microsoft Excel^®^ and GraphPad Prism 5^®^. Differences were considered statistically significant at the p<0.05 level. For each sample, 4 images were taken with a 4X, 10X or 20x magnification depending on the experimental design. Cell quantification and analysis was performed using ImageJ^®^. Phalloidin staining (F-actin) images of single cells were analysed for filament coherency using the Orientationβ plugin for ImageJ ^46^.

## Supplementary Figures

**Figure S1.**
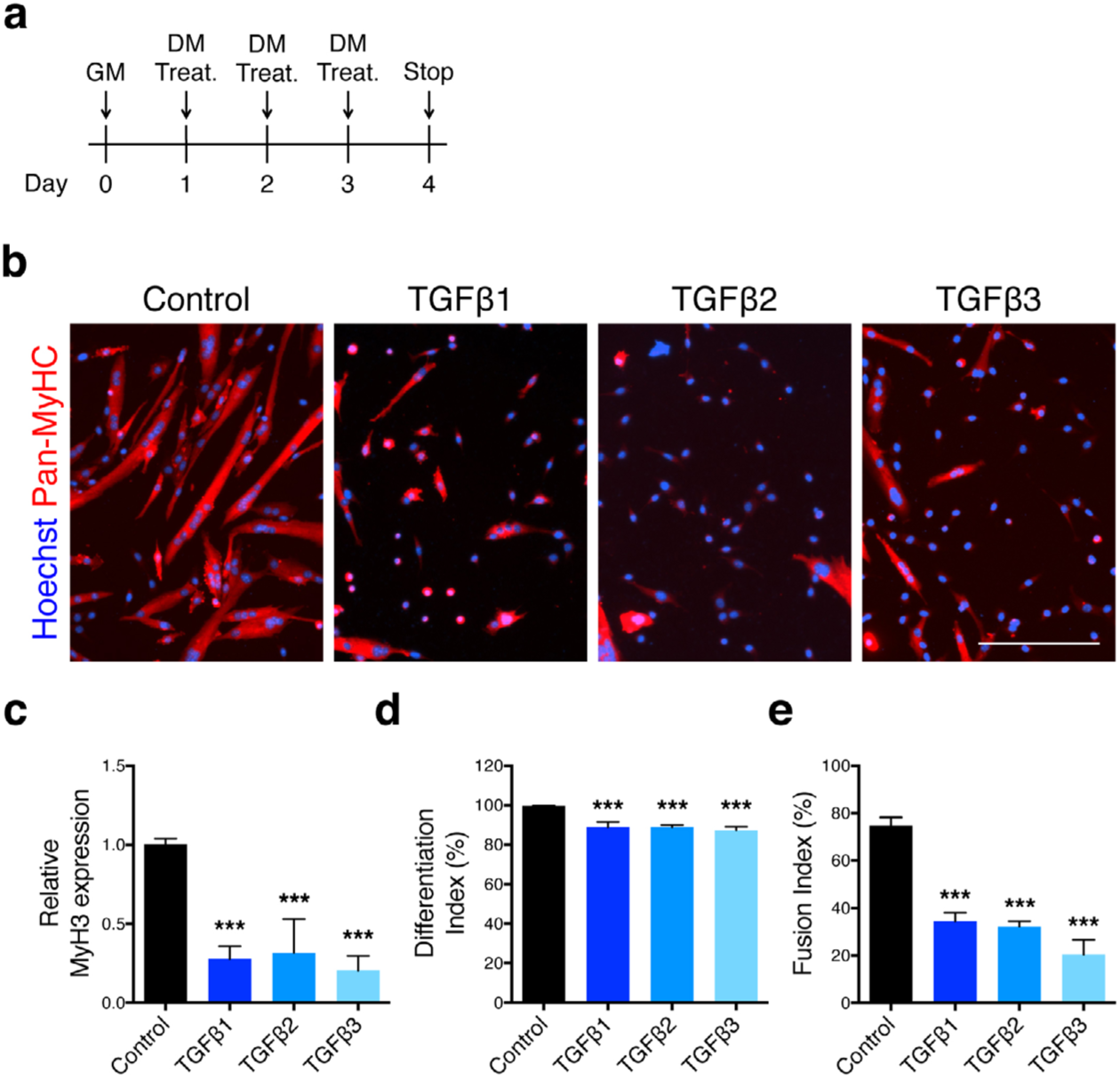
TGFβ signaling effects on *in vitro* myogenic differentiation. **a,** Experimental scheme. Primary myoblasts were induced to differentiate in medium containing TGFβ recombinant proteins. **b,** Immunofluorescent staining for Pan-MyHC of 3-days differentiated myotubes. **c,** qRT-PCR analysis for Myh3 (Embryonic Myosin Heavy Chain) transcript expression by of 3-days differentiated primary myoblasts indicates that stimulation of the pathway downregulates its expression compared to the control. **d,** Percentage of Pan-MyHC-expressing cells of 3-days differentiated primary myoblasts. **e,** Fusion index of 3-days differentiated primary myoblasts shows that TGFβ stimulation inhibits fusion. Scale bars: **c,** 1000μm. Data are presented as mean ± s.e.m. from at least three independent experiments. N.D.=Not significant, compared with Control (Unpaired two-tailed Student’s t-test).

**Figure S2.**
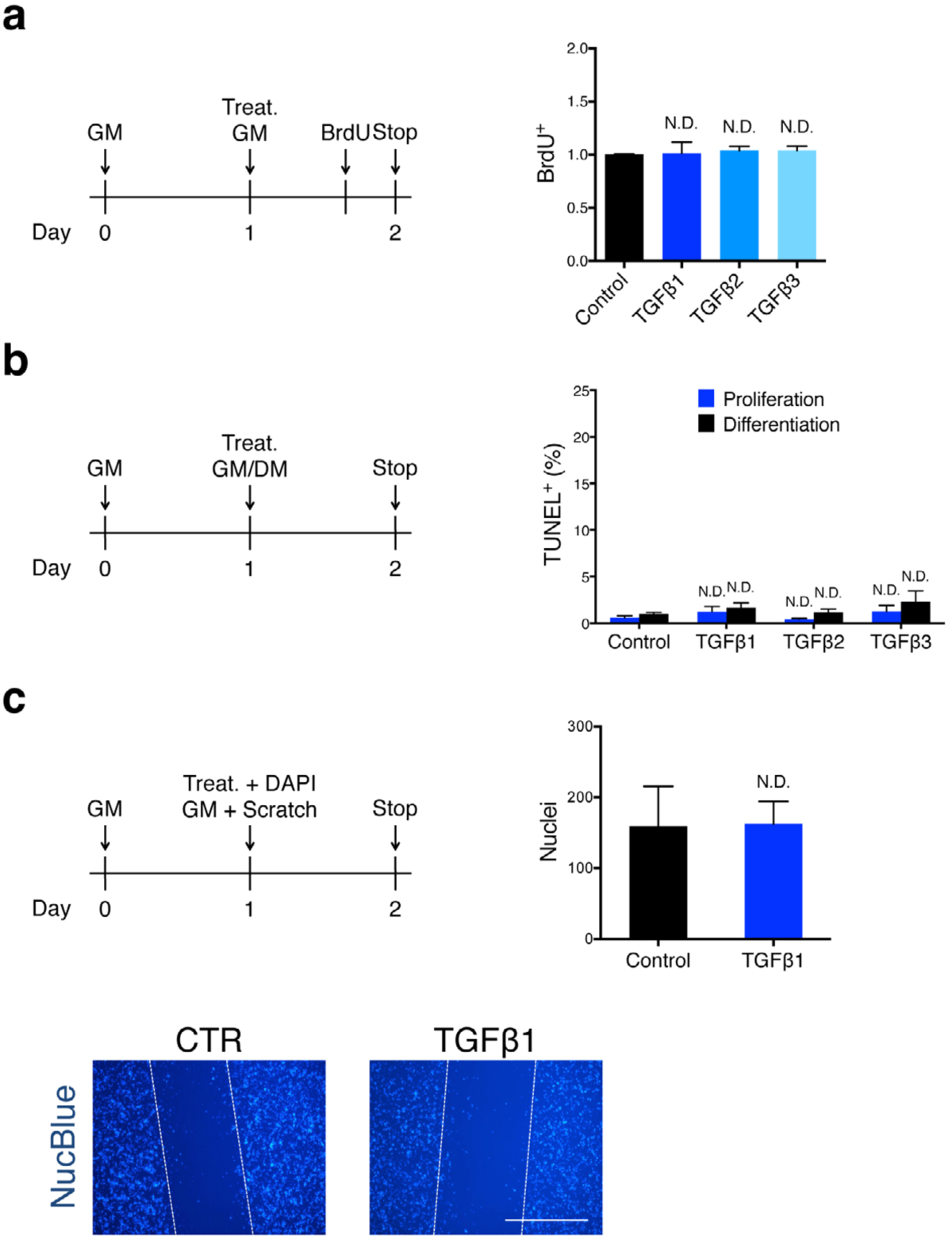
TGFβ signaling does not affect myoblast proliferation, death and motility. **a,** Primary myoblasts were treated with TGFβ1, 2 or 3 for 24h and incubated with BrdU for the last 40 minutes before fixation. 0uantification of BrdU^+^ cells shows no differences. **b,** Primary myoblasts were treated with TGFβ1, 2 or 3 for 24h in proliferating or differentiating conditions. TUNEL^+^ cells were quantified, and no particular death rates were detected. **c,** Primary myoblasts were treated with TGFβ1, 2 or 3 for 24h in proliferating condition. When treated, cell layer was scratched and washed with PBS. Scratch-wound images were taken after 24 hours of treatment. Cells were stained with NucBlue. The quantification of nuclei within the scratch-wound reveals that motility is not affected. Scale bars: **b,** 200μm. Data are presented as mean ± s.e.m. from at least three independent experiments. \*\*\**P*<0.001, compared with Control (Unpaired two-tailed Student’s t-test).

**Figure S3.**
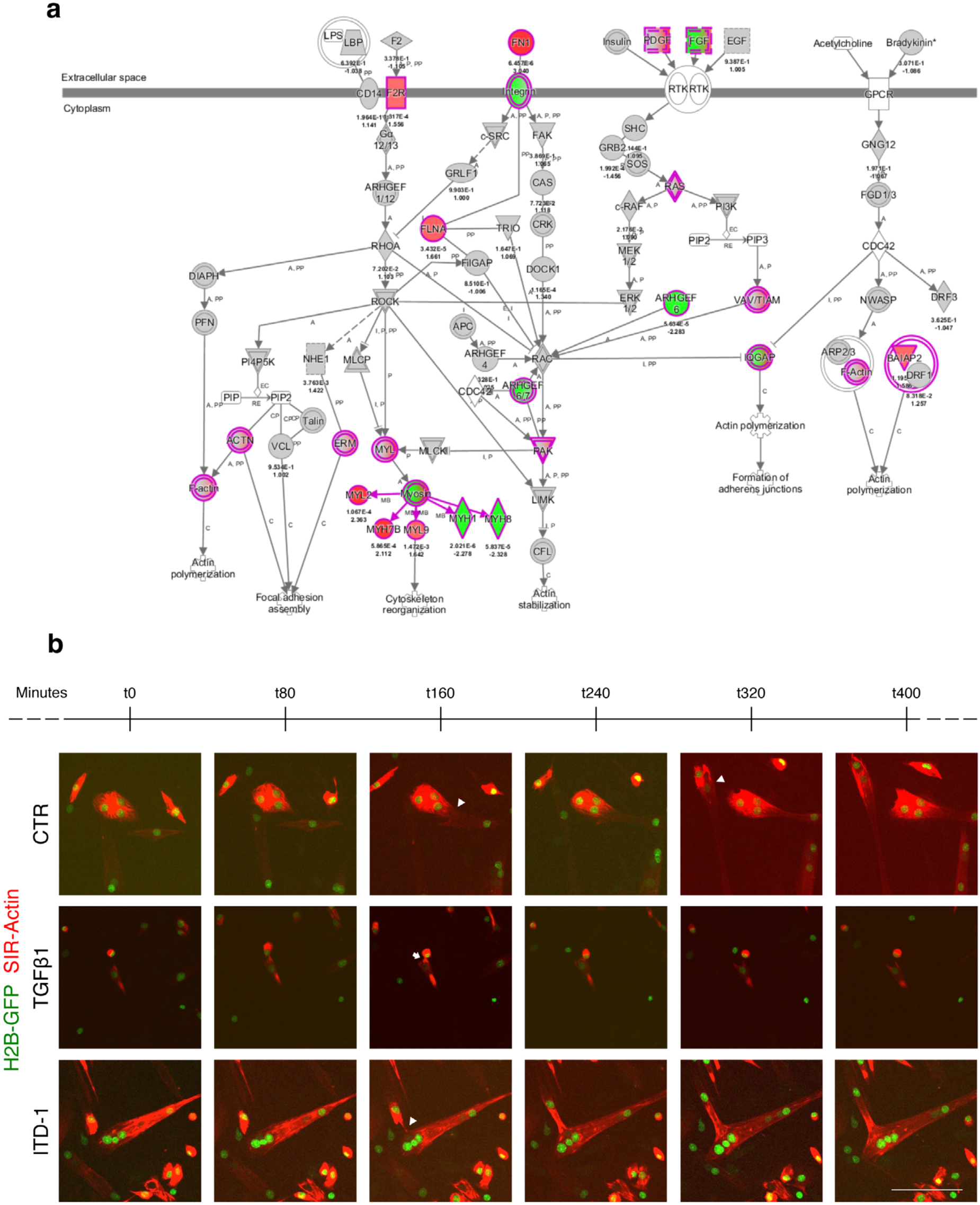
Effects of TGFβ cascade on Actin signaling pathway. **a,** Representative scheme of the Actin signaling pathway genes modulated by TGFβ cascade. **b,** Live-imaging frames of pre-differentiated myocytes expressing H2B-GFP and stained with SiR-Actin. Arrowheads indicate fusion events, arrow depicts cell-cell interaction. In control condition, fusion occurs linearly, while ITD-1 treatment allows perpendicular fusion. On the other hand, TGFβ1 allows cell-cell interactions, but blocks fusion. Scale bars: **b,** 200μm. Data are presented as mean ± s.e.m. from at least three independent experiments.

**Figure S4.**
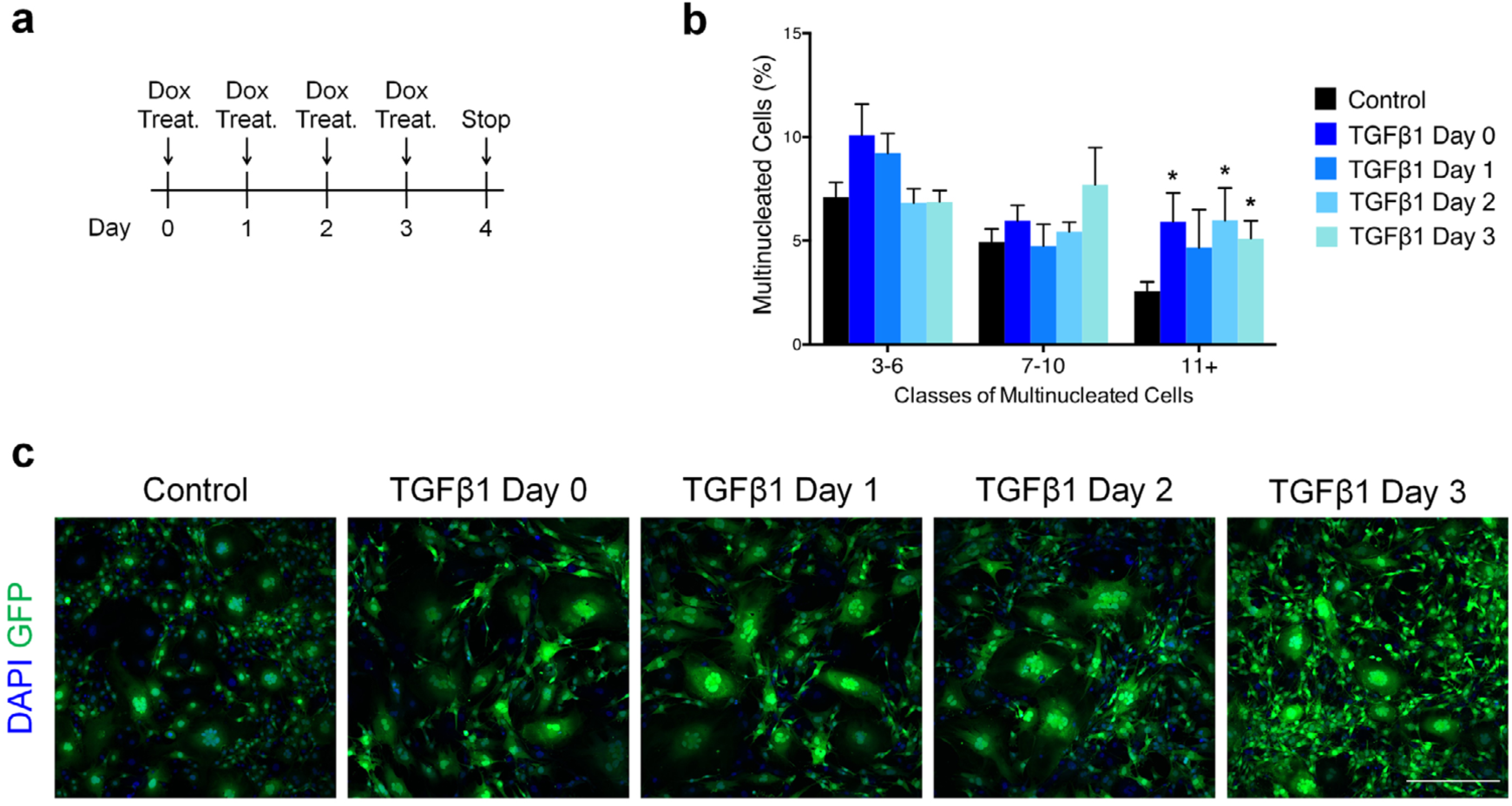
TGFβ signaling exerts its effect independently from Myomaker and Myomerger. **a,** Experimental scheme. A dox-Inducible Myomaker- and Myomerger-expressing fibroblasts were used to test TGFβ1 effect on cell-cell fusion. Dox was administrated at day 0 and refreshed every day, while TGFβ1 either at day 0, 1, 2 or 3. **b,** Aggregation index of 4-days Myomaker and Myomerger-expressing fibroblasts showing no significant reduction of the fusion process when TGFβ1 is administrated compared to the control. **c,** GFP-myomaker-infected fibroblasts, transduced with dox-inducible myomerger, were visualized with fluorescent microscopy. TGFβ stimulation does not reduce the fusion process. Scale bars: **c,** 400μm. Data are presented as mean ± s.e.m. from at least three independent experiments. \**P*<0.05 compared with Control (Unpaired two-tailed Student’s t-test).

